# Memory’s gatekeeper: the role of PFC in the encoding of familiar events

**DOI:** 10.1101/2024.02.01.578356

**Authors:** Inês C. Guerreiro, Claudia Clopath

## Abstract

Theoretical models conventionally portray the consolidation of memories as a slow process that unfolds during sleep. According to the classical Complementary Learning Systems (CLS) theory (as presented in J. McClelland et al., 1995), the hippocampus (HPC) rapidly changes its connectivity during wakefulness to encode ongoing events and create memory ensembles that are later transferred to the prefrontal cortex (PFC) during sleep. However, recent experimental studies challenge this notion by showing that new information consistent with prior knowledge can be rapidly consolidated in PFC during wakefulness, and that PFC lesions disrupt the encoding of familiar events in the HPC. These results challenge the widely accepted view that consolidation is a slow process that unfolds during sleep and highlight the role of PFC during the initial stages of memory encoding. The contributions of the PFC to memory encoding have therefore largely been overlooked. Moreover, most theoretical frameworks assume random and uncorrelated patterns representing memories, disregarding the correlations between our experiences. To address this shortcomings, we developed a HPC-PFC network model that simulates interactions between the HPC and PFC during the encoding of a memory (awake stage), and subsequent consolidation (sleeping stage) to examine the contributions of each region to the consolidation of novel and familiar memories. Our results show that the PFC network uses stored memory “schemas” consolidated during previous experiences to identify inputs that evoke familiar patterns of activity, quickly integrated it in its network, and gate which components are encoded in the HPC. More specifically, the PFC uses GABAergic long-range projections to inhibit HPC neurons representing input components correlated with a previously stored memory “schema”, eliciting sparse hippocampal activity during exposure to familiar events, as it has been experimentally observed.

## Introduction

A fundamental question in memory research is how new memories in a labile state are transformed into more robust and permanent memories. A widely accepted framework posits that during learning, the hippocampus (HPC) encodes new information, enabling rapid acquisition of ongoing events without interfering with existing neocortical knowledge. This process is followed by hippocampal replay, during sleep, which effectively ‘teaches’ the recently acquired information to the prefrontal cortex (PFC) (Marr, 1971; J. McClelland et al., 1995).

While extensive experimental and theoretical work support this view (Alvarez and Squire, 1994; Diekelmann and Born, 2010; Goode et al., 2020; Klinzing et al., 2019; Marr, 1971; Milner and Penfield, 1955; Nadel et al., 2007; Rolls, 1996; Sekeres et al., 2018; Singh et al., 2022; Squire, 1986; Teyler and DiScenna, 1986), recent experimental studies highlight the involvement of the PFC during the initial stages of learning (Dash et al., 2004; Kitamura et al., 2017; Miyawaki and Mizuseki, 2022), suggesting that it might have a broader role than previously thought, extending beyond offline memory consolidation. Notably, lesions and pharmacological inactivation of PFC disrupts the learning of spatial (Churchwell et al., 2010; DeVito and Eichenbaum, 2010; Kyd and Bilkey, 2003), and familiar memory tasks, i.e. tasks that are small variations of previously learned ones (DeVito et al., 2010; Guise, 2017; Tse et al., 2011). While the role of PFC in memory encoding (during wakefulness) remains elusive, experimental studies indicate that prior consolidation of a PFC associative memory “schema” – pre-existing network of connected neocortical representations (van Kesteren et al., 2018) - enables rapid learning of congruent (i.e. familiar) information (Tse et al., 2007; van Kesteren et al., 2013). Interestingly, encoding of congruent information has been correlated with a decrease in hippocampal activity (Cohen et al., 2017; Karlsson and Frank, 2008; Lee et al., 2020; van Kesteren et al., 2013). Furthermore, Guise and Shapiro have shown that mPFC inactivation reduced hippocampal pattern separation of overlapping hippocampal representations (Guise, 2017), a phenomenon typically ascribed to processes supported by the hippocampal neural circuitry, in particular in dentate gyrus (Leutgeb et al., 2007; Neunuebel and Knierim, 2014; O’Reilly and McClelland, 1994; Sakon and Suzuki, 2019; Treves and Rolls, 1994; Yassa and Stark, 2011). Taken together, these results emphasize the need to reevaluate the mechanisms by which functions HPC-PFC interactions support memory processing.

In this study, we propose a computational model of interacting HPC and PFC networks to examine how the PFC modulates hippocampal activity during the encoding of familiar versus novel memories, and how prior knowledge consolidated in PFC influences the learning rate of new information. In addition to excitatory HPC-to-PFC projections considered in previous memory models (for example, Alvarez and Squire, 1994; Marr, 1971; Singh et al., 2022; Squire, 1986), our model includes long-range GABAergic PFC-to-HPC connections, as recently reported in Malik et al., 2022. Our computational model simulates interactions between the hippocampus and prefrontal cortex during the encoding of a memory (Awake stage) and subsequent consolidation (Sleeping stage). We begin by presenting the naive network with a new pattern representing a to-be-learnt memory. After confirming that the pattern has been successfully encoded in the HPC and then consolidated into the PFC, we evaluated the responses of both the HPC and PFC when presented with an additional input pattern which is familiar (i.e., overlapping). We then examined the distinct contributions of HPC-to-PFC and PFC-to-HPC interactions. The simulations capture the rapid encoding of information consistent with prior knowledge described experimentally in Tse et al., 2007, and propose a circuit mechanism through which the PFC uses pre-existing memory schemas to guide the integration of new information into the HPC-PFC network. Furthermore, our modeling work shows that the PFC creates sparse representations of familiar inputs in the HPC, enabling pattern separation.

## Results

We first aim to investigate how the mechanisms underlying the long-term storage of novel and familiar information differ and the specific roles of the hippocampus and PFC in these processes over time. To that end, we begin by storing a pattern A into an untrained (i.e. naive) HPC-PFC network. Once pattern A is consolidated in the PFC, the network receives a pattern B which can either overlap (familiar) or not (novel) with the neural representation of pattern A. In our model, the HPC and PFC are described as recurrent neural networks with plastic Hebbian all-to-all intra-regional connections (with a higher learning rate for HPC than for PF), and fixed one-to-one inter-regional connections. In addition to excitatory HPC-to-PFC connections and a projection of the external inputs onto the HPC considered in conventional memory models, we implement inhibitory PFC-to-HPC connections, as reported in Malik et al., 2022. Based on anatomical studies (Golmayo et al., 2003), we also include a projection of the external inputs, i.e. of the pattern to be encoded, onto the PFC.

### Encoding and consolidation of a memory in a naive neural network

We start by storing a pattern A in the HPC-PFC network that can be used as a reference to compare and classify future incoming inputs as novel or familiar. Storing pattern A involves submitting the network to the awake stage, when it receives pattern A, followed by the sleeping stage.

Starting from a naive state (HPC and PFC connectivity: *W*_*HPC*_ = *W*_*PFC*_ = 0; HPC and PFC initial activity: *x*_*HPC*_(*t*0) = *x*_*PFC*_(*t*0) = 0), the HPC and PFC receive a pattern A to be encoded (awake stage; Figure 1 (a1)). At the end of the awake stage, we found strong recurrent connections within the HPC network amongst neurons activated by pattern A (Figure 1 (a2)), indicating that a memory trace of pattern A is encoded in HPC. The PFC connectivity, on the other hand, remained unchanged. This is due to the fact that the PFC learning rate is smaller than the one of HPC.

**Figure 1:**
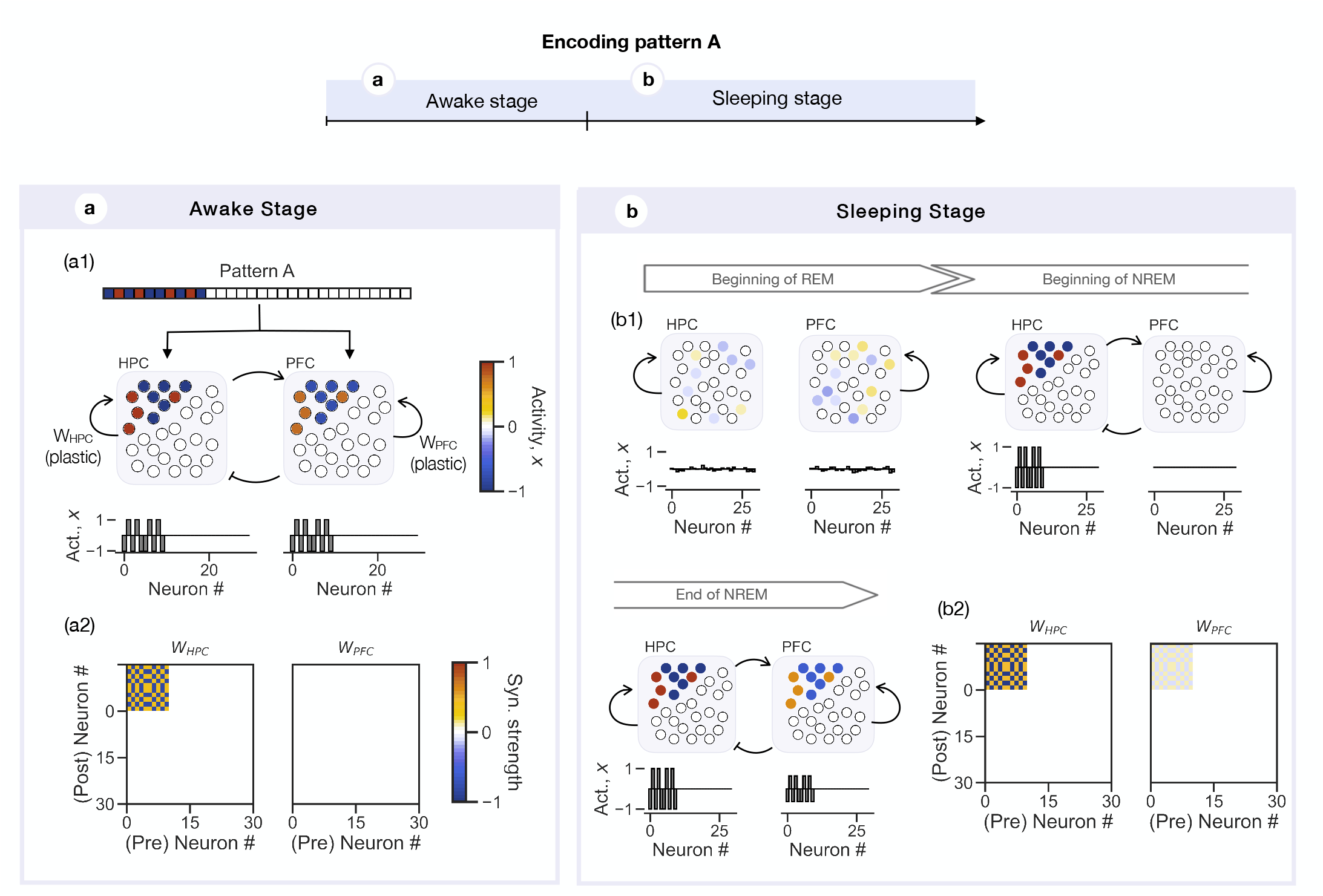
Encoding and consolidation of a memory in a naive neural network. Top panel: The HPC-PFC network encodes a pattern A. The network goes trough two stages: an awake stage, where the network receives pattern A, and a sleeping stage, where the network evolves autonomously according to its intrinsic dynamics. **(a)** During the awake state, the hippocampus (HPC) and prefrontal cortex (PFC) network receive a pattern A represented by ones (1; red entries) and minus ones (−1; blue entries), targeting the first 10 neurons in the HPC and PFC network. The recurrent connections are plastic. The two regions are coupled through fixed one-to-one HPC-to-PFC excitatory, and PFC-to-HPC inhibitory connections (*W*_*HPC−PFC*_ = 0.5, *W*_*PFC−HPC*_ = *−* 1). Each circle represents a neuron of the network, with the color and the height of the corresponding bars representing its activity (top panel and bottom panel, respectively). At the end of the awake stage, the HPC and PFC show the same pattern of activation, with the HPC units more strongly activated (a1), and the HPC connectivity has formed an engram of pattern A (a2). **(b)** The sleeping stage is characterized by a REM phase, when the two regions are uncoupled (*W*_*HPC−PFC*_ = *W*_*PFC−HPC*_ = 0), and a NREM phase, when the two regions are coupled through excitatory HPC-to-PFC and inhibitory PFC-to-HPC connections (*W*_*HPC−PFC*_ = 1, *W*_*PFC−HPC*_ = *−* 1). During the sleeping stage, the system cycles through the REM and NREM phases seven times. Every time the network enters the REM phase, the HPC and PFC networks are reset to a noisy random state, from which it evolves according to its intrisinc dynamics. In this case, the HPC converges to memory pattern A and the PFC decays to its naive state (b1; first sleep cyle). At the end of the sleeping stage, the memory engram A is consolidated in the PFC connectivity (b2).

Once the HPC has formed a memory trace of input A, the HPC-PFC network enters the sleeping stage, where it cycles through a rapid eye movement (REM) (uncoupled phase; *W*_*PFC−HPC*_ = *W*_*HPC−PFC*_ = 0) and a non-rapid eye movement (NREM) (coupled phase; *W*_*PFC−HPC*_ = *−*1, *W*_*HPC−PFC*_ = 1) seven times (Figure 1 (b1)). Every time the network enters the REM stage, the HPC and PFC neurons are reset to a random noisy state, and the system evolves autonomously according to its intrinsic dynamics. This enables replay of recently acquired information during sleep, which is believed to facilitate learning (Cairney et al., 2015; Diekelmann and Born, 2010; Rasch and Born, 2013). In our case, the HPC network will converge to the memory engram A, which means that all neurons encoding memory pattern A will become activated in a similar fashion as during wakefulness. At the end of the sleeping stage, PFC activity reflects the neural representation of pattern A (Figure 1 (b1), bottom panel), and its connectivity resembles the hippocampal memory engram A (Figure 1 (b2)), although with weaker connectivity weights. Nonetheless, pattern A is consolidated in PFC, as it was confirmed by testing the PFC ability to recall pattern A upon partial activation of the its neural ensemble (Figure S1).

### Consolidation of novel memory pattern relies on hippocampal replay during sleep

Once pattern A has been consolidated in PFC, we set out to examine the HPC-PFC network’s behavior when receiving a novel pattern B (0% overlap with pattern A). We consider that a long time has passed since consolidation of pattern A, and that while the PFC network retained the memory trace of A encoded in its connectivity, hippocampal connectivity decayed back to its initial naive state (*W*_*HPC*_ = 0). In other word, the HPC does not hold any information about pattern A; it has forgotten it.

When presented with pattern B, the hippocampal and PFC neural ensembles targeted by the novel pattern were strongly activated (Figure 2 a1)). At the end of the awake stage, an engram of pattern B was encoded in the HPC connectivity (Figure 2 a2)). In contrast, at this stage, the PFC did not incorporate the novel input into its network. Only after going through the sleeping stage was pattern B consolidated in the PFC (Figure2 b2)).

**Figure 2:**
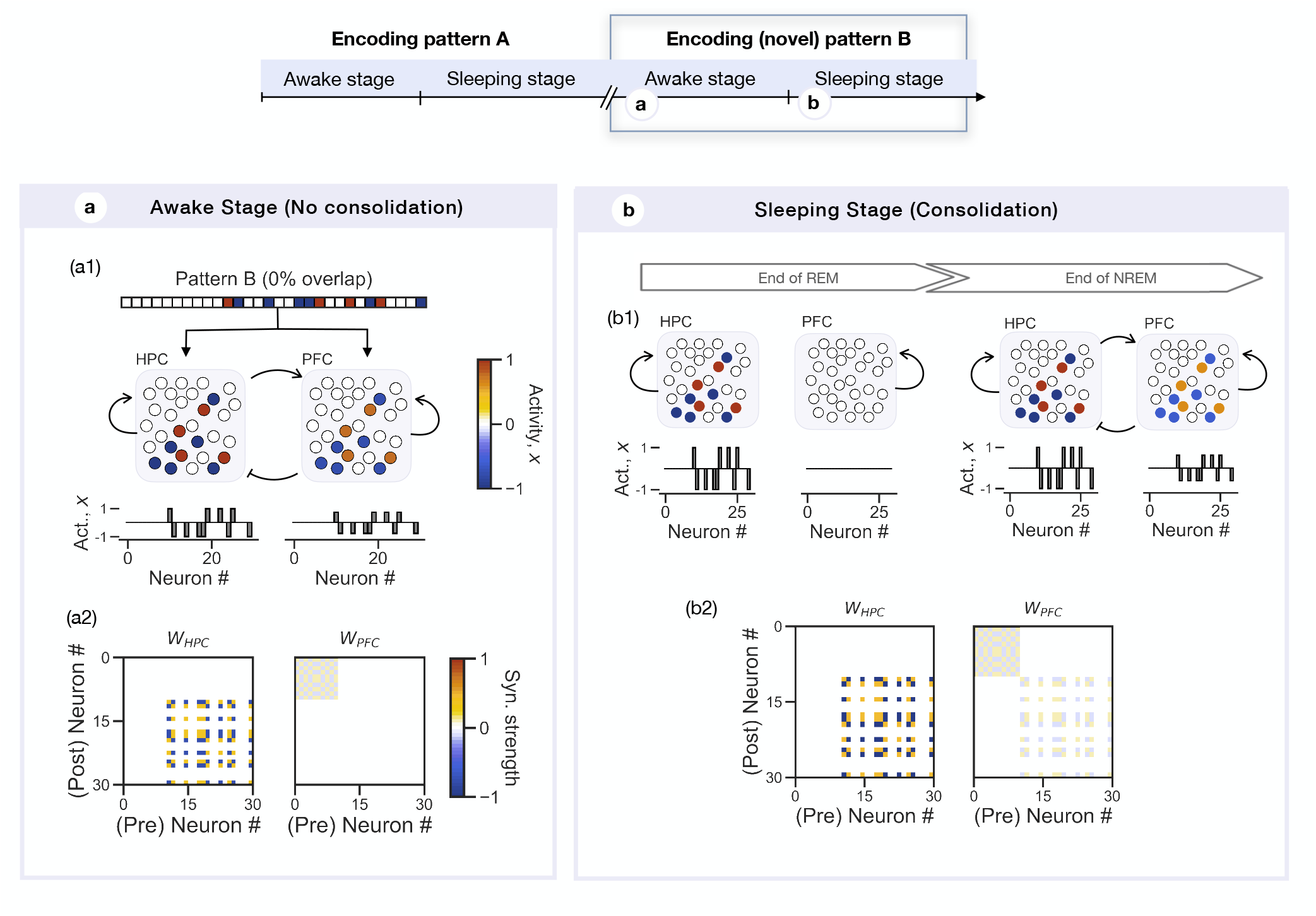
Consolidation of novel memory pattern relies on hippocampal replay during sleep. Top panel: Hippocampal and prefrontal cortex activity are analyzed during the awake stage, when the network receives a pattern B whose representation does not overlap with the previously consolidated pattern A (overlap 0%), meaning that it targets a different neural ensemble. Encoding of pattern B happens after consolidation of pattern A in the PFC, and decay of its engram in HPC, i.e. when the recurrent hippocampal connectivity is back to its naive state. **(a)** During the awake state, the hippocampus (HPC) and prefrontal cortex (PFC) network receive a pattern B targeting 10 HPC and PFC units uncorrelated with the units encoding for pattern A (0% overlap). At the end of the awake stage, the HPC and PFC show the same pattern of activation, with the HPC units more strongly activated. **(b)** The hippocampal network has encoded pattern B in its connectivity, *W*_*HPC*_, forming a memory engram B. The PFC connectivity, *W*_*PFC*_ remains unaltered, i.e. it only encodes the memory engram A.

Interestingly, following the reset of the HPC and PFC to a noisy state in the beginning of the REM state, the PFC does not evolve towards the previously consolidated pattern A state of activity (Figure 2 b1), End of REM). This is due to the connectivity of the memory engram of pattern A, which is not strong enough to drive the PFC network to it and the network decays back to its resting state (*x*_*PFC*_ = 0). This suggests that replay during sleep is mainly driven by hippocampal activity, which in this case will evolve towards the novel pattern B.

Our modeling results suggest that incongruent knowledge previously consolidated in the PFC does not influence the mechanisms of long-term storage of novel information - the HPC-PFC network exhibit a similar behavior to the naive case. In other words, consolidation of novel information in PFC relies on offline (during sleep) replay of hippocampal activity learned during wakefulness. These results are consistent with the idea that the there is a fast-learning system in HPC that quickly stores information online, which can be replayed to a slower learning system in PFC (Marr, 1971; J. McClelland et al., 1995).

### Familiar pattern is quickly stored during wakefulness

We next sought to examine the effects of previous knowledge (pattern A) on consolidation of congruent (i.e. familiar) information (pattern B with 90% overlap with A).

Contrarily to what was observed for the case of a novel pattern, when presented with a familiar input, the PFC quickly integrated its uncorrelated components (highlighted in Figure 3 a2)) in its connectivity, suggesting a rapid consolidation of pattern B during the awake stage. The HPC, on the other hand, did not form a memory trace of pattern B (Figure 3 a2), left panel). During the sleeping stage, there was no replay of pattern B, and the HPC and PFC connectivity remained the same as at the end of the awake stage (Figure 3 b2)).

**Figure 3:**
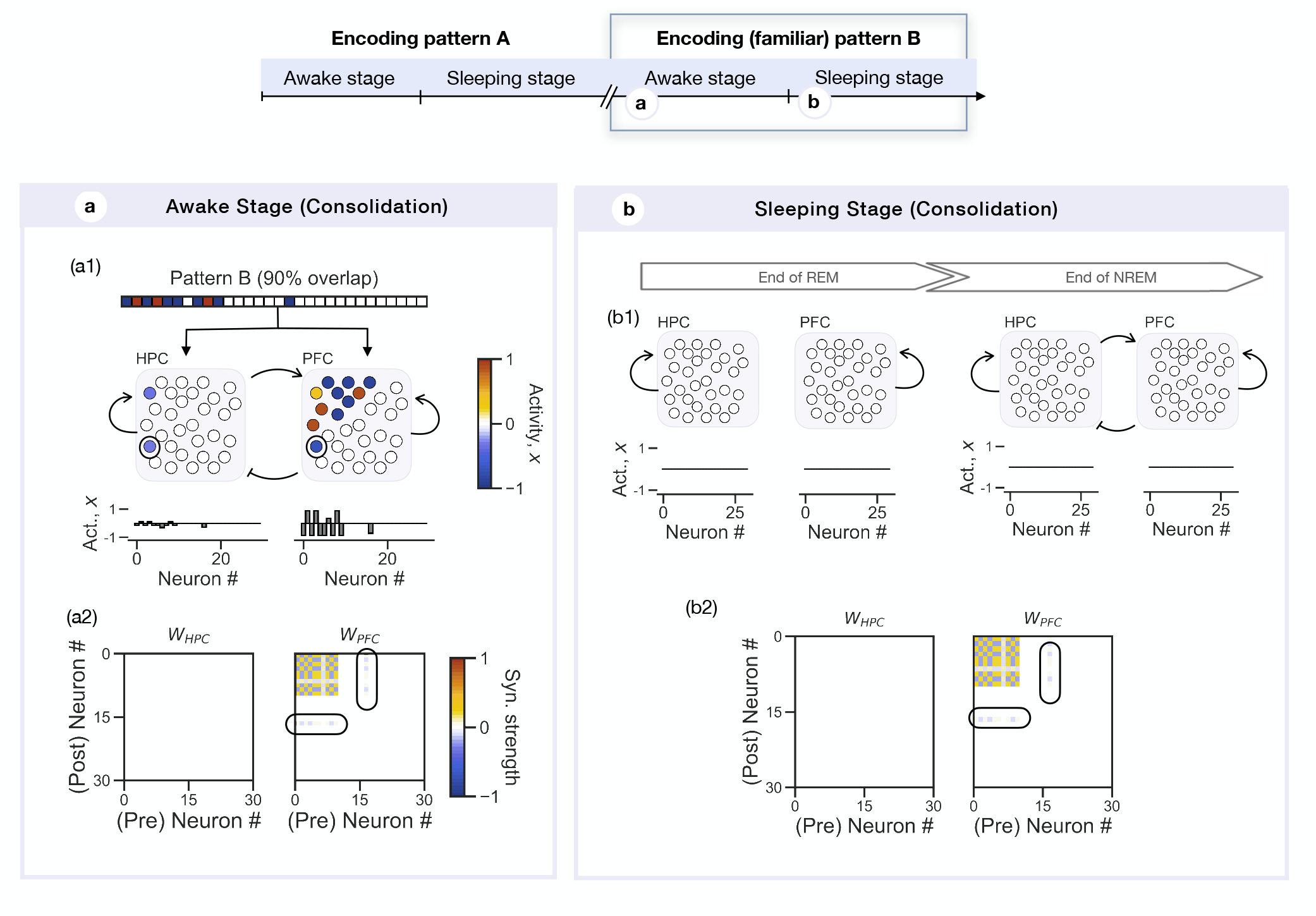
Familiar pattern is quickly stored during wakefulness. Top panel: Hippocampal and prefrontal cortex activity are analyzed during the awake stage, when the network receives a pattern B whose representation overlaps with previously consolidated pattern A by 90%. Encoding of pattern B happens after consolidation of pattern A in the PFC, and decay of its engram in HPC. **(a)** The HPC network shows sparse activity, while the PFC units targeted by input B are strongly activated. **(b)** At the end of the awake stage, the HPC network connectivity remains unaltered (i.e. in its naive state). The PFC network, on the other hand, has integrated the uncorrelated components of pattern B in its connectivity with the memory engram A. The circle highlights the non-overlapping, i.e. uncorrelated, components of pattern B.

Interestingly, we see that HPC neurons targeted by the familiar pattern B were weakly activated, apart from the ones representing the input components uncorrelated with pattern A (highlighted units in Figure 3 (a)), contrasting to what was observed in the case of a novel pattern. We hypothesize that this is directly modulated by inhibition from strongly activated PFC neurons.

Given the substantial overlap (90%) between the representations of patterns A and B, activation of the pattern B neural ensemble in the PFC drives the system into the attractor state formed during the consolidation of pattern A. As a result, PFC neurons that were part of the engram of A and were targeted by B became highly activated, driven by both external inputs and potentiated recurrent intrinsic connections (see Figure S2). This high activity led to rapid plasticity. The hippocampal neurons representing the correlated portion were suppressed by strongly activated PFC cells. The strong activation of pattern B components in PFC paired with activation of uncorrelated components driven by external and hippocampal excitatory inputs (highlighted components in Figure 3) led to changes in its connectivity, *W*_*PFC*_, to encode the familiar pattern B. Furthermore, common features between patterns A and B were reinforced in PFC connectivity.

### Differential roles for HPC and PFC in the encoding of familiar inputs

We next sought to examine the contributions of the inter-regional connections to the rapid and sparse hippocampal activity observed during encoding of a familiar pattern. For that, we repeated the simulations where we present the HPC-PFC network with the same familiar pattern B, but set either the excitatory HPC-to-PFC or the inhibitory PFC-to-HPC to zero. Suppression of the HPC-to-PFC excitatory connections (*W*_*HPC−PFC*_ = 0, *W*_*HPC−PFC*_ = *−*1) impaired encoding of the familiar pattern B in PFC during wakefulness (Figure 4 (a)), while suppressing PFC-to-HPC inhibitory connections (*W*_*HPC−PFC*_ = 0.5, *W*_*HPC−PFC*_ =) abolished the sparse hippocampal activity previously observed, with the HPC encoding for the full representation of pattern B (Figure 4 (b)). These results suggest that the bidirectional inter-regional HPC-PFC connections contribute in distinct ways to the behavior of the HPC-PFC network during encoding of familiar events. In particular, hippocampal excitatory inputs facilitate rapid encoding of familiar patterns in PFC, while PFC inhibition mediates HPC sparse activity.

**Figure 4:**
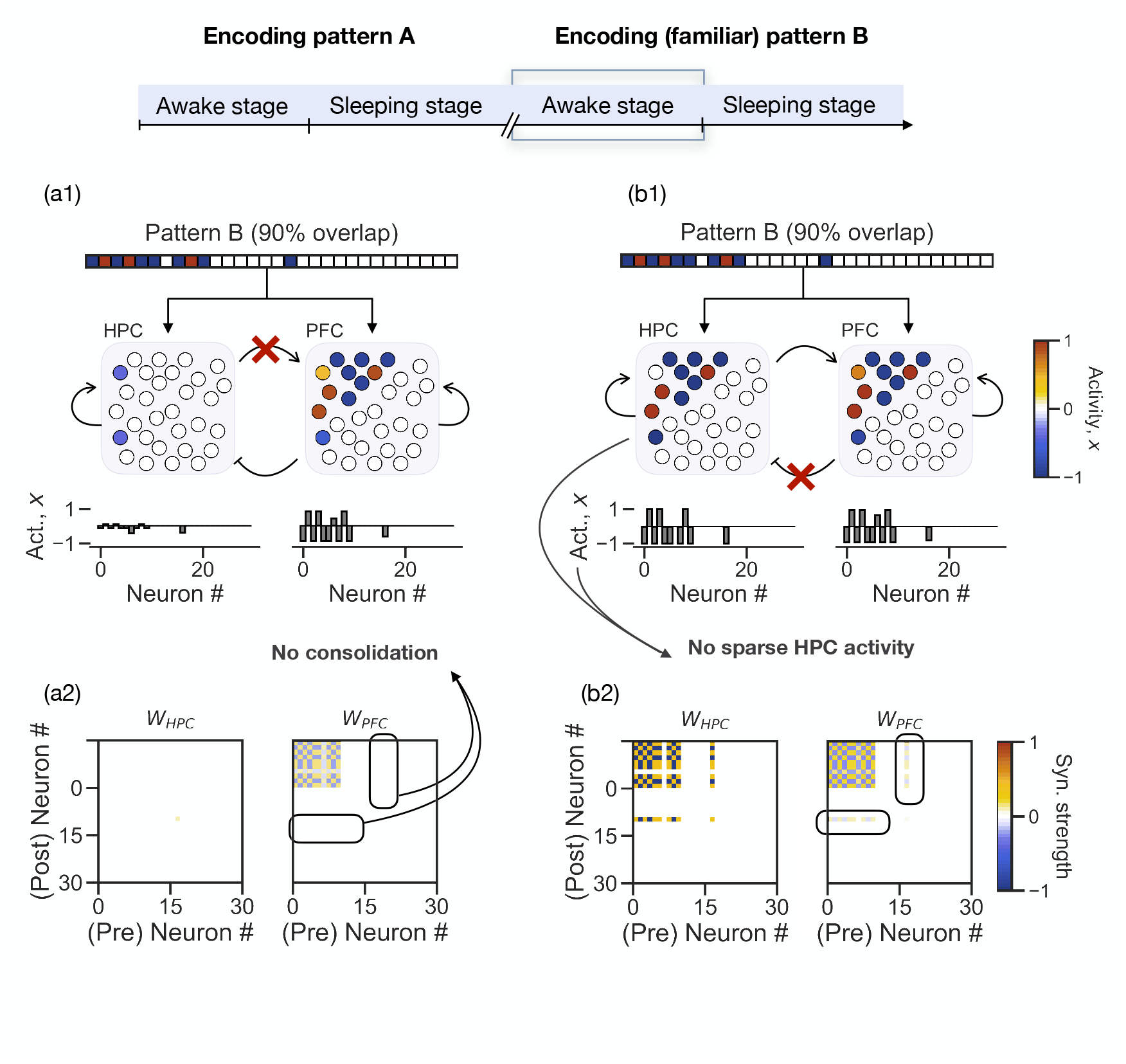
Differential roles for HPC and PFC in the encoding of familiar inputs. **(a)** The HPC-PFC network receives an input B overlapping with memory A by 90%. Coupling between the two regions is only mediated by inhibitory PFC projections (*W*_*HPC−PFC*_ = 0, *W*_*PFC−HPC*_ = *−* 1). At the end of the awake stage, the PFC network did not encode for the uncorrelated components of input B in its connectivity, indicating that the input was not consolidated. **(b)** The HPC-PFC network receives an input B overlapping with memory A by 90%. Coupling between the two regions is only mediated by excitatory HPC projections (*W*_*HPC−PFC*_ = 1, *W*_*PFC−HPC*_ = 0). At the end of the awake stage, both the hippocampal and PFC network have encoded input B in its connectivity. However, we no longer have the sparse hippocampal activity observed during the encoding of familiar memories (Cohen et al., 2017; Karlsson and Frank, 2008; Lee et al., 2020).

### Examining influence of degree of familiarity of new information in PFC plasticity and HPC-PFC network activity

We next sought to examine how these results generalize to degrees of overlap between patterns A and B that range from 0 to 90% (instead of considering just these two extreme cases). More specifically, we wanted to know if there is a well-defined threshold at which the PFC network identifies an incoming pattern as familiar, triggering rapid consolidation and hippocampal sparse activity, or whether it is a smooth and graded process where PFC memory traces of pattern B become stronger the bigger the degree of overlap, and gradually inhibit more HPC neurons. For that, we quantified changes in connectivity of PFC neurons targeted by pattern B for different degrees of overlap (0-90% with increments of 10%) during the awake stage, and estimated the mean HPC and PFC activity (Figures 5 (a) and (b), respectively).

**Figure 5:**
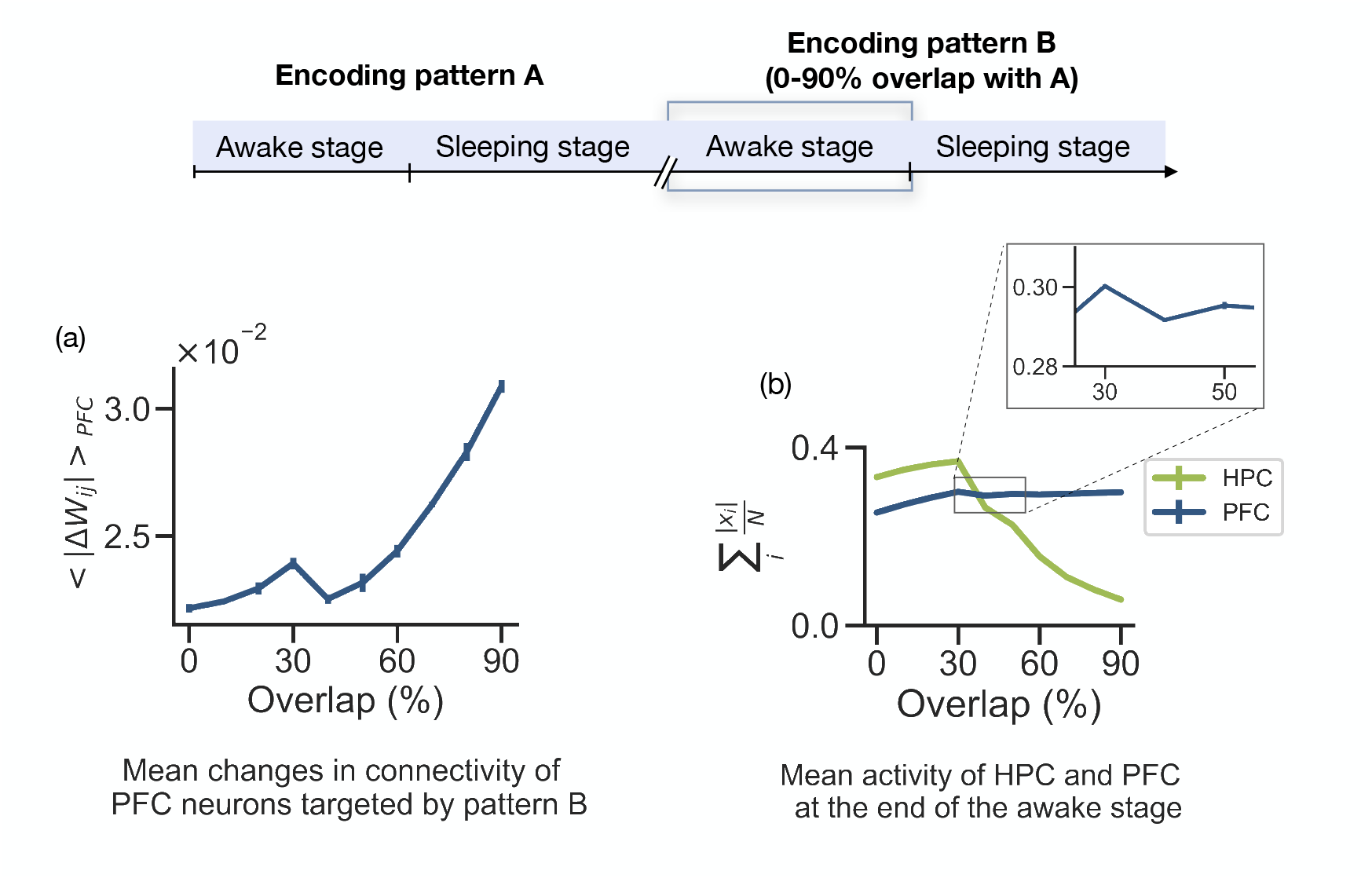
Examining influence of degree of familiarity of new information in PFC plasticity and HPC-PFC network activity. Top panel: HPC and PFC activity and changes in connectivity are analyzed during the awake stage, when the network receives a pattern B whose representation overlaps with previously consolidated pattern A by 0-90%. **(a)** Mean absolute changes of the PFC connections between neurons encoding pattern B, estimated at the end of the awake stage, for patterns B overlapping by 0-90% with A. For each degree of overlap, we considered 10 randomly generated patterns B, and the average of the mean connectivity changes obtained for each pattern. **(b)** Mean absoulte activity of all the HPC and PFC neurons (green and blue line, respectively). Once more, for each degree of overlap we considered 10 randomly generated patterns B.

We found that while there is a tendency to have a stronger memory trace of pattern B (i.e. bigger changes in PFC connectivity encoding for pattern B) the bigger the overlap with the stored pattern A, we observe a decrease connectivity changes when pattern B overlaps with A by 40% and 50% compared with the case of 30%. The same propensity appears in the mean PFC activity, which shows a slight increase with the degree of overlap, except for an overlap of 40 and 50% (Figure 5 (b), inset). This indicates that, in the framework here considered, the PFC is able to adopt a fast (slow) speed of consolidation when the incoming information is clearly familiar (novel), i.e. it overlaps by 90% (0%) with previous knowledge. However, if an incoming pattern overlaps by 40-50%, the network shows an ambiguous behaviour. These results align with previous indications that memories tend to be stronger when the encoded information either aligns with our previous knowledge or is completely novel (Alonso et al., 2020). Surprisingly, we found that there is a clear threshold for which hippocampal activity dramatically decreases (Figure 5 b)). This result can be explained if we consider a form of a race between the HPC and PFC regions are to form a memory trace of an incoming pattern B. During the awake stage, both the HPC and PFC receive the a pattern at the same time and with the same characteristics. As they do, the activity of the HPC and PFC neurons encoding the pattern starts to increase until it reaches a point (*x*_*i*_ = 0.4) where they start to strengthen their intra-regional connections with co-activated neurons. If the PFC neurons reach a level of activation comparable with the level of activation of HPC neurons they target before HPC has the chance to significantly change its connections with other neurons representing pattern B, then their activity is suppressed by PFC.

### Familiar inputs are linked in PFC, whereas unfamiliar stimuli exhibit pattern separation

To examine how are familiar inputs integrated with consistent knowledge in PFC, we tested the ability of PFC to recall memory engram A and B (memory linking), or just memory engram B (pattern separation) at the end of the awake stage, when, as we have seen in the previous section, familiar inputs are consolidated, and at the end of the sleeping stage, when the HPC-PFC network replays the patterns encoded during wakefulness (Figure **??**). To that end, we examined the response of engram A and B neurons to activation of a subset of engram B cells. If all neurons of engram B were activated, but not engram A, we have that the two memories are stored independently (pattern separation). If both engrams A and B were activated to activation of engram B, then the two memories were linked together (memory linking). For simplicity, we start by analyzing the two extreme cases: when pattern B overlaps by 10% (novel) and 90% (familiar) with A. If pattern B is novel (only overlaps by 10% with the representation of A) activating 90% of engram B cells at the end of the awake stage (Before sleep test) won’t prompt the recall of the either the pattern B or A in PFC (Figure 6 (a1)). However, if pattern B is familiar (overlaps by 90% with A) activation of solely 30 engram B units recall not only the full pattern B, but also pattern A, indicating that at the end of the awake stage, the two memories are linked in PFC (Figure 6 (a2)). When performing the same test at the end of the sleeping stage, we now get that activation of 90% of engram cells of a novel pattern B causes recall of B, without recall of A (Figure 6 (b1)), indicating pattern separation of the two overlapping representation. For a familiar pattern B, the network shows the same performance as after the awake stage, i.e. activation of 30% engram B cells recalls both engrams A and B (Figure 6 (b2)).

**Figure 6:**
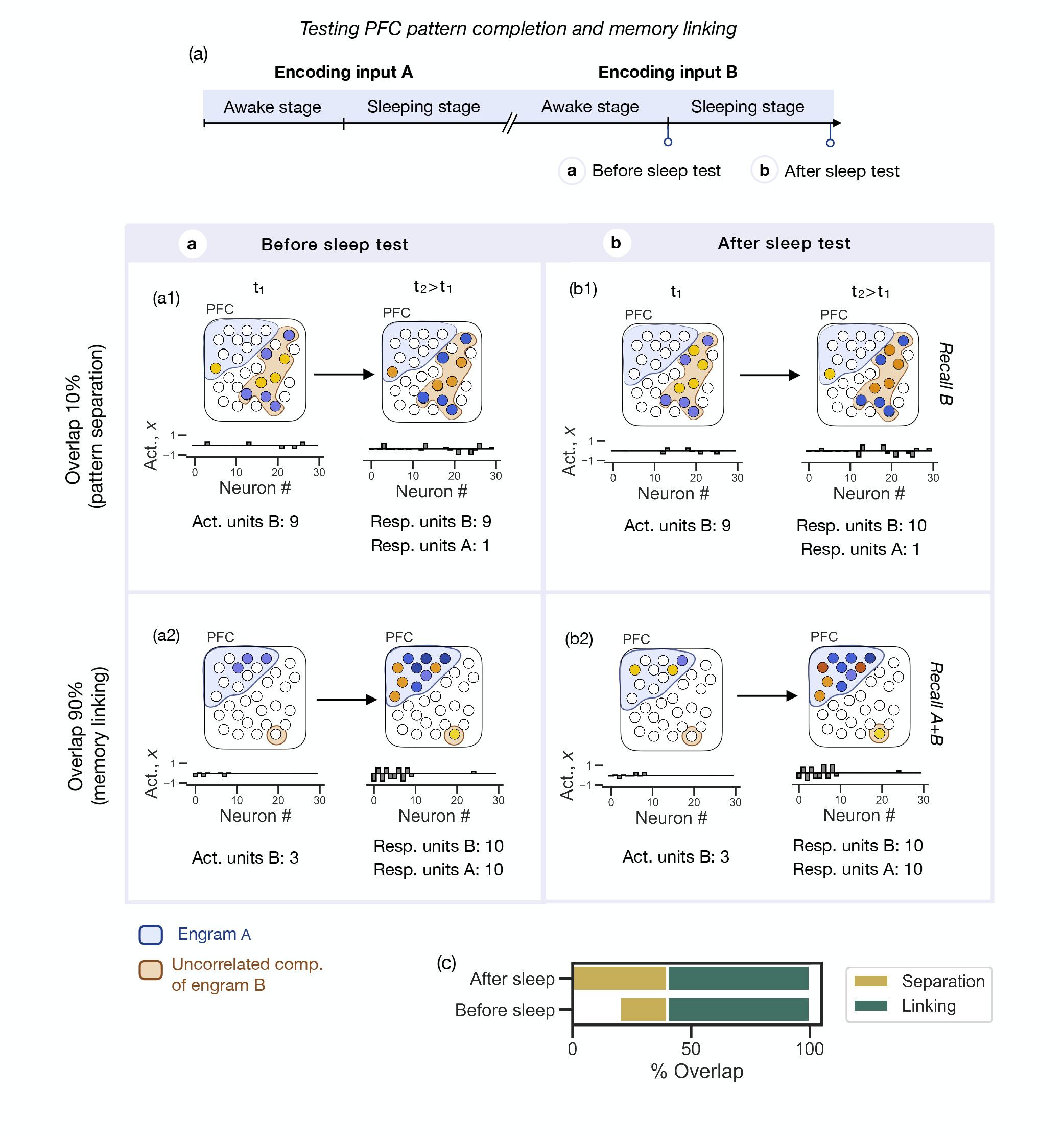
Familiar inputs are linked in PFC whereas unfamiliar stimuli exhibit pattern separation. Top panel: Testing PFC ability to perform pattern separation and memory linking at the end of the awake stage (a) and at the end of the sleeping stage (b). **(a)** Testing the PFC ability to recall pattern B and pattern A at the end of the awake stage. If pattern B overlaps by 10% with pattern A, the PFC won’t be able to recall engram B or engram A (a1). The PFC pattern of activation 90 time steps and 7600 time steps after activating 9 out of 10 engram B units is the same (left and right panel, respectively). If pattern B overlaps by 90% with pattern A, activating 3 engram B units results in the recall of both engram A and engram B, indicating that the two patterns are linked (i.e. activation of engram A plus the uncorrelated components of engram B; a2). **(b)** Testing the PFC ability to recall pattern B and pattern A at the end of the sleeping stage. If pattern B overlaps by 10% with pattern A, activating a subset of engram B units results in recall of engram B but not engram A, indicating pattern separation (b1). If pattern B overlaps by 90%, activation of a subset of engram B units recalls engram A and B, similarly to what was observed at the end of the awake stage (b2). **(c)** Classifying pattern separation and memory linking at the end of the awake stage (Before sleep) and at the end of the sleeping stage (After sleep) for patterns B overlapping by 0-90% with pattern A. Pattern separation is defined as recall of engram B without recall of A. Familiar inputs (overlap *>*40%) are encoded in PFC during awake stage and are linked to previously consolidated overlapping representations. Novel inputs are separated and consolidated during sleep.

Overall, our results show that there is a familiarity threshold (40% overlap) above which the PFC network rapidly integrates incoming input with previously previously stored congruent patterns, linked both representations. On the hand, inconsistent information (30% overlap) relies on HPC-PFC activity replay during sleep to be consolidated in the PFC network without interfering with previously consolidated knowledge (Figures 6 (c) and S5). Note that, according to our model, there is an intermediate result (overlap between 30 and 40%) for which incoming inputs are familiar enough to evoke recurrent activation of a correlated schema and rapidly consolidate this information with the uncorrelated components of the input, but the overlapping schema is not big enough to recruit the full engram of the previously stored consistent pattern. This means that such inputs are rapidly consolidated during wakefulness, but are not linked to consistent patterns.

We also note that the more familiar an incoming input is (the bigger the overlap with stored information), the easier it is to recall it. For example, considering the cases where pattern B overlaps by 50 and 90% with A, both patterns are consolidated in PFC during wakefulness and linked to pattern A. However, with a 90% overlap we only need to activate 30% of engram B cells to recall it, while recall for the case of a 50% overlap requires the activation of 90% engram B cells (Figure S5).

Considering that, for a certain degree of overlap, the neurons representing the the correlated components of a pattern B are randomly selected, different patterns are going to have the same degree of overlap with pattern A. While examining how different patterns (10) with a same degree of overlap affect the results, we found that we have approximately consistent results (same as for Figure 6 c)) except for pattern with a degree of overlap of 40% with stored information. When testing recall of pattern B with a 40% overlap with A at the end of the awake stage, for certain patterns the PFC could not recall it (see Figure S6 for examples a pattern B with 40% overlap resulting in a successful and unsuccessful recall). This aligns with previous results showing that there is a break in the general tendency of increased changes in the PFC connectivity during wakefulness with the degree of overlap at 40% (see Figure 5 a)), reinforcing the idea that the ease in which ‘in-between’ events (in this case, with a familiarity degree of 40%) are recalled is inconsistent. This ambiguous behavior is particularly accentuated during the awake stage of encoding, with results stabilizing during the sleep.

Altogether, our model proposes that familiar inputs are linked in PFC to congruent information during wakefulness, while inconsistent information relies on HPC and PFC replay during sleep to be consolidated without interfering with stored knowledge.

## Discussion

In the wild, adult animals rarely encounter new information in isolation. Their experiences are usually linked to what they’ve encountered before. Previous research shows that past experiences affect how new memories are processed (Bartlett, 1932; Harlow, 1949). For instance, consider how easily you grasp new information related to your field compared to unrelated information. However, most memory studies overlook the impact of previously acquired knowledge in experiments. In recent work, Tse et al., 2007, 2011 showed that new associations consisted with a previously consolidated PFC schema quickly becomes HPC-independent, suggesting rapid PFC consolidation. Consequent theoretical work by McClelland has shown that a Rumelhart network is able to integrate new information into existing consistent knowledge (J. L. McClelland, 2013). However, such network use a back-propagation learning rule and are not biologically plausible. Thus, the neural and circuits mechanisms underlying the rapid consolidation of congruent information remain elusive.

In this work, we hypothesize that schemas stored in PFC promote rapid integration of congruent events without sleep by enabling strong activation of engram cells representing the overlapping representation of the stored and new events. According to our model, rapid consolidation relies on activation of PFC neurons by an external and hippocampal input. By adopting an Hebbian learning rule, where changes in synaptic strength depend on the level of activation of the pre- and post-synaptic units, we can overcome the slow learning rates characteristic of the PFC by strongly activating the interacting neurons that form a memory engram. A familiar external input drives the PFC network to the closest attractor state, previously form during the consolidation of a similar pattern and characterized by strong recurrent connections. Neurons activated by both the external input and strong recurrent connections, i.e. neurons representing the correlated components of the familiar input, will be activated strongly enough to quickly strengthen their connections with neurons representing the uncorrelated components. In other words, the ‘novel part’ of a familiar input will readily be incorporated with the stored schema. If, on the other hand, the circuit receives a novel input, the PFC network won’t converge to an attractor state, and activation of the targeted neurons will be solely due to the action of the external input. Neurons representing the novel input will be weakly activated, and won’t be able to strengthen their synapses and form an engram. In this case, the HPC-PFC network needs to go through the sleeping stage, where the HPC repeatedly re-activates the neural ensembles representing the novel input, in order to consolidate it. It is important to note that, in this case, our model is in line with conventional accounts of memory consolidation mechanisms (for example, J. McClelland et al., 1995).

Our model captures several important experimental findings; namely, the quick consolidation of events consistent with prior knowledge, that is disrupted by removal of the hippocampus (Tse et al., 2007, 2011). Our results grant a broader role to the PFC in the encoding of events than classically considered. Besides hypothesizing the PFC potential to consolidate memory events without hippocampal intervention, we also propose its potential role in supporting HPC by modulating its activity during the encoding phase. When incoming information is congruent with previously acquired knowledge, the PFC quickly incorporates its uncorrelated components with a pre-existing overlapping schema in its network through strengthening of its cortico-cortical functional connections, and inhibits the hippocampal activity encoding for the correlated components of the new input.

Many studies have suggested that PFC exerts top-down control over information processing in the HPC (Benoit et al., 2015; Eichenbaum, 2017; Malik et al., 2022; Preston and Eichenbaum, 2013; Rajasethupathy et al., 2015). Here, motivated by recent anatomical studies reporting long-range GABAergic projections from PFC to HPC Malik et al., 2022, we propose a network mechanism through which the PFC exerts top-down inhibition over hippocampal activity during the encoding of familiar events. We propose that this mechanism can be responsible for the sparse activity seen in the hippocampus during the encoding of familiar events. A large body of work has focused on hippocampal contributions to pattern separation (Leutgeb et al., 2007; Neunuebel and Knierim, 2014; O’Reilly and McClelland, 1994; Sakon and Suzuki, 2019; Treves and Rolls, 1994; Yassa and Stark, 2011). However, converging evidence suggests that pattern separation is supported by a network of brain regions (Amer and Davachi, 2023). Here, we predict that PFC can contribute to hippocampal pattern separation by suppressing hippocampal activity encoding information already consolidated in PFC. Such a mechanism could also promote increased memory capacity of the HPC.

In summary, our modeling work suggests the following: 1) rapid encoding of consistent information in PFC during wakefulness is mediated by hippocampal inputs and strong activation of a congruent PFC schema; 2) inconsistent information relies on hippocampal replay during sleep to be consolidated; 3) familiar information is integrated in PFC into a congruent schema whereas novel information undergoes pattern separation, producing little interference with memories already stored in PFC; 4) GABAergic PFC-to-HPC projections induce sparse hippocampal activity during the encoding of familiar events, enabling pattern separation.

## Methods

### Multi-region Recurrent Neural Network

The hippocampus and prefrontal cortex regions are modeled as rate-based recurrent networks. Each region is composed of N=30 units with all-to-all connections. The dynamics of each unit *x*_*i*_ is described by the following equation:

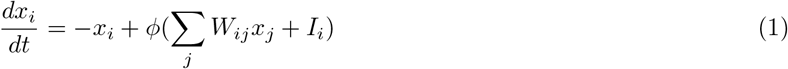

where *ϕ* is a nonlinearity applied to the total input each unit receives. IT is modeled as *ϕ*(*s*) = *tanh*(*x*) for hippocampal units, and *ϕ*(*x*) = *tanh*(0.5*x*) for prefrontal cortex units, reflecting differences in responsiveness of neurons in both regions (Ruggiero et al., 2021). More specifically, the PFC units exhibit a more gradual response compared to HPC. *W*_*ij*_ is the synaptic weight between the pre- and post-synaptic units, *j* and *i*, respectively, and *I*_*i*_ the external input. We considered that both regions receive identical external inputs. Patterns of ones (1) and minus ones (−1) describe the memory we want to store into the network. We consider inputs represented by neural ensembles with 10 units. In other words, an external input is composed of 10 entries of values 1 or -1, and the remaining 20 entries are zero. We assume that different memories are represented by different neural ensembles of the same size. We will use the term ‘familiar input’ to refer to an external input that is represented by a subset of neurons that overlaps with the representation of a previously stored input.

Intra-regional connections are dynamic and change according to the following standard Hebbian learning rule:

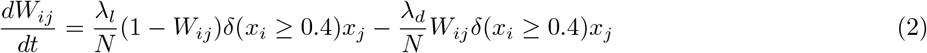

where *λ*_*l*_ is the learning rate (*λ*_*l*_(*HPC*)=0.45, *λ*_*l*_(*PFC*)=0.06), and *λ*_*d*_ the decay rate (*λ*_*d*_(*HPC*)=0.55, *λ*_*d*_(*PFC*)=0). A gating mechanism has been introduced to the learning rule, contingent on the activity of the neurons involved. When the activities of both pre- and post-synaptic neurons, *x*_*j*_ and *x*_*i*_, exceed a threshold of 0.4, the standard Hebbian term contributes to the adjustment of synaptic strengths, aiming to reinforce connections that are correlated with neuronal activity. This gating mechanism enables the learning rule to account for situations where neuronal activity is insufficient to drive synaptic potentiation, allowing for a more biologically realistic representation of learning dynamics in the network.

Inter-regional connections are mediated through one-to-one excitatory HPC-to-PFC connections, and reciprocate inhibitory PFC-to-HPC connections. Note that, these connections are not of the same nature. Excitatory functional connections from HPC to PFC are mainly modulated by intermediate regions (for example, the thalamus Tomé et al., 2022). In this case, the strength of the connections *W*_*HPC−PFC*_ reflect the coupling strength of hippocampal and PFC oscillatory activity. On the other hand, the inhibitory PFC to HPC connections represent long-range GABAergic projections, i.e. direct connections from the PFC into HPC (Malik et al., 2022). The value of *W*_*PFC−HPC*_ is representative of the conductance of GABA receptors in HPC neurons.

For simulations, the differential equations were solved using Euler’s method with a time step Δ*t* = 0.01 (*a*.*u*.).

### Simulating awake and sleep

During the awake stage, the HPC and PFC receive an external input, *I*. The two regions are coupled through one-to-one excitatory HPC-to-PFC connections (*W*_*HPC−PFC*_ = 0.5), and one-to-one inhibitory PFC-to-HPC projections (*W*_*HPC−PFC*_ = *−*1). The awake stage has a duration of 7600 time steps.

Following the awake stage, the model enters the sleeping period. The model has a rapid eye movement (REM) sleep stage, where the two regions are uncoupled and evolve autonomously according to their intrinsic dynamics, and a non-rapid eye movement (NREM) sleep stage, where dynamics between the hippocampus and PFC are tightly coupled (reference). During sleep, coupling is modeled by setting *W*_*HPC−PFC*_ = 1 and *W*_*PFC−HPC*_ = *−*1. This set up is inspired by previous studies suggesting that during NREM sleep, the coupling between the HPC and PFC tends to be relatively strong, while in REM sleep, the coupling between the HPC and PFC is generally weaker Cairney et al., 2015; Diekelmann and Born, 2010. We find that alternating between REM and NREM sleep stages 7 times facilitates consolidation of hippocampal-dependent memories into the PFC network, which has a small learning rate. Each time the model enters the REM stage, the hippocampal and PFC neurons are reset to a noisy state Rasch and Born, 2013. The state of each neuron is randomly drawn from a normal distribution with standard deviation 0.1 and mean 0.

The REM and NREM stages have a duration of 9000 and 900 time steps, respectively.

### Model simulations

Starting the network from a naive state, i.e. with hippocampal and prefrontal cortex connectivities set to zero (*W*_*HPC*_ = 0, *W*_*PFC*_ = 0), the network receives a pattern A to encode. Pattern A is represented by 10 HPC and PFC units; in other words, it targets the first 10 neurons of each region (Pattern A = [-1 1 -1 1 -1 -1 1 -1 1 -1 0 … 0]). Storing pattern A involves submitting the network to the awake stage, when it receives pattern A, followed by the sleeping stage, instead of being set up directly in the PFC network. We do so to insure that the strength which pattern A is encoded in the PFC connectivity is a result of the natural awake-sleep cycle to avoid introducing a bias in simulations that follow.

After going through the awake and sleeping stage, we verify that pattern A was successfully consolidated in the PFC network (see). We then assume that a long time has passed (let’s say, months) and that, while the PFC network retained the memory trace of A in its connectivity, the hippocampal network decayed back to its naive state. At this point, the network receives an input B whose representation overlaps with pattern A.

Pattern B is generated by randomly selecting a fraction *f* of the first 10 entries of pattern A to remain unaltered (i.e. the entries with values 1 and -1) and the rest of the initial 10 entries (10-*f*) are set to zero. The parameter *f* reflects the extent of overlap between patterns A and B. For instance, to generate a pattern B with an 80% overlap with A, 8 out of the 10 first entries in pattern A remain the same. To ensure that any disparities in activity observed in the HPC and PFC result exclusively from their interactions, and not because input B targets a different number of units, out of the 20 zero entries of pattern A, we randomly select (10-*f*) entries to change to 1 or -1 (also in a random independent way).

### Testing pattern completion, pattern separation and memory linking

To evaluate the PFC network’s ability to perform pattern completion, we assessed the responsiveness of neurons to the activation of a subset of engram cells. A neuron, denoted as *i*, was deemed responsive if after 8000 time steps |*x*_*i*_| *>* 0.2, while activity below this threshold was considered noise. Successful pattern completion refers to the network’s capacity to reactivate all cells within a memory engram when a subset is activated. This implies the consolidation of the memory within the network. The degree of consolidation is determined by the size of the subset required for pattern completion, which we refer to as the ‘Recall threshold’. The smaller the subset of cells needed to achieve pattern completion, the stronger the consolidation of the memory. To examine pattern separation and memory linking, we tested the network’s ability to perform pattern completion of engram B when a subset of engram B neurons were activated, as well as the pattern completion of engrams A and B when a subset of engram B cells were activated. The successful pattern completion of both engrams when only a subset of engram B is activated indicates the linking of both memories. On the other hand, pattern completion of engram B without engram A signifies pattern separation. To ensure the accuracy of our findings, we assessed the responsiveness of the total number of cells forming an engram to the activation of subsets of different sizes. The subset of cells to be activated was randomly selected. For each subset size, we repeated the analysis 50 times, activating a different subset of neurons each time.

### Sparsity index

In this study, two distinct approaches are utilized to quantify and analyze sparsity. Firstly, we investigate the amplitude and position of density distribution peaks in the HPC and PFC activity. A higher peak centered around zero indicates sparser activity, suggesting that a majority of neurons within the network exhibit zero activity. To visualize these distributions, we employ kernel density estimation (KDE) plots. Secondly, we calculate a sparsity index. This index is determined by computing the average absolute activity across all neurons in the network, represented as 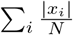. A lower value of the sparsity index suggests sparser activity. To examine the sparsity dynamics during the encoding of various types of information, we conduct the analysis for a range of inputs that have different degrees of overlap (0% to 90%) with a previously stored memory.

## Supplementary Figures

**Figure S1:**
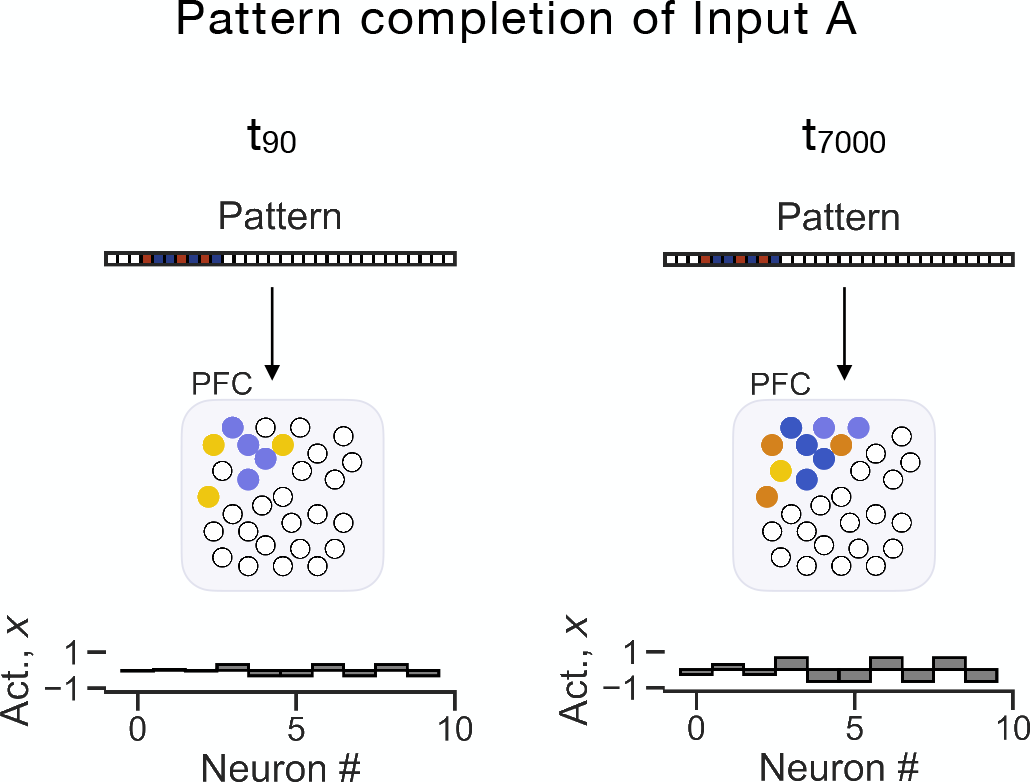
Pattern A is consolidated in PFC. PFC activity was examined at 90 and 7000 time steps, when activated by a pattern partially overlapping with pattern A. A pattern targeting 7 out of 10 engram A units, enables the recall of the full engram A pattern of activity, demonstrating the PFC ability to perform pattern completion of pattern A.

**Figure S2:**
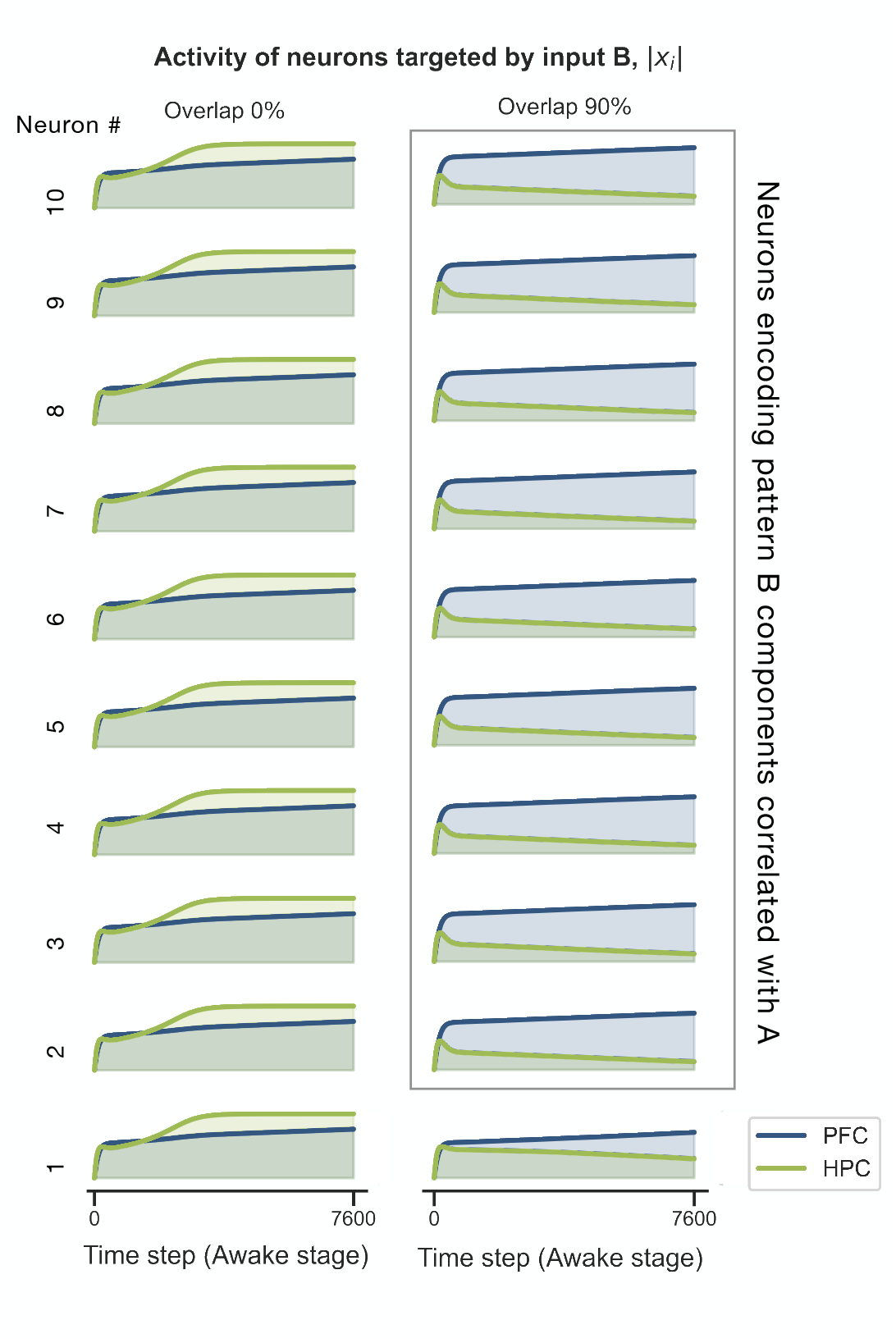
Familiar patterns enable strong activation of PFC neurons and suppression of hippocampal activity. Left panel: a novel pattern will evoke weak PFC activity, and the activity of HPC neurons is not suppressed. Right panel: a familiar input evokes strong PFC activity that suppresses HPC neurons representing the correlated components of pattern B with A.

**Figure S3:**
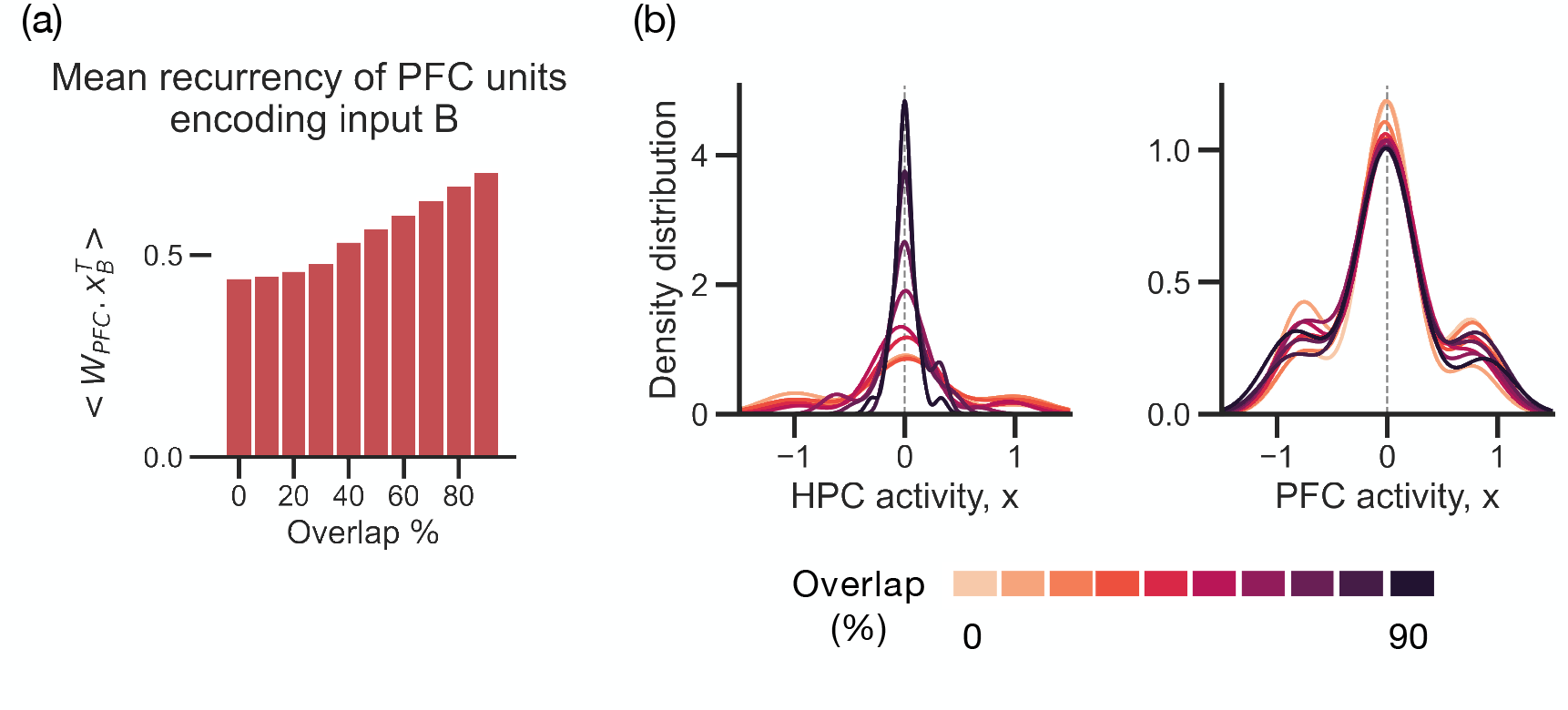
Familiar inputs elicit strong recurrent activity in PFC and sparse hippocampal activity. **(a)** Mean recurrent input the PFC units representing pattern B receive, *Wx*^*T*^, at 800 a.u. (time units), during the awake stage. The bigger the overlap between patterns B and A, the stronger the activation of PFC units due to intrinsic recurrent inputs. **(b)** Density distribution of hippocampal and prefrontal cortex activity upon receiving a pattern B (left and right panel, respectively). Patterns B sharing 0-90% of common features with input A are analyzed. The bigger the overlap between inputs A and B, the weaker is hippocampal activity, and the stronger PFC activity.

**Figure S4:**
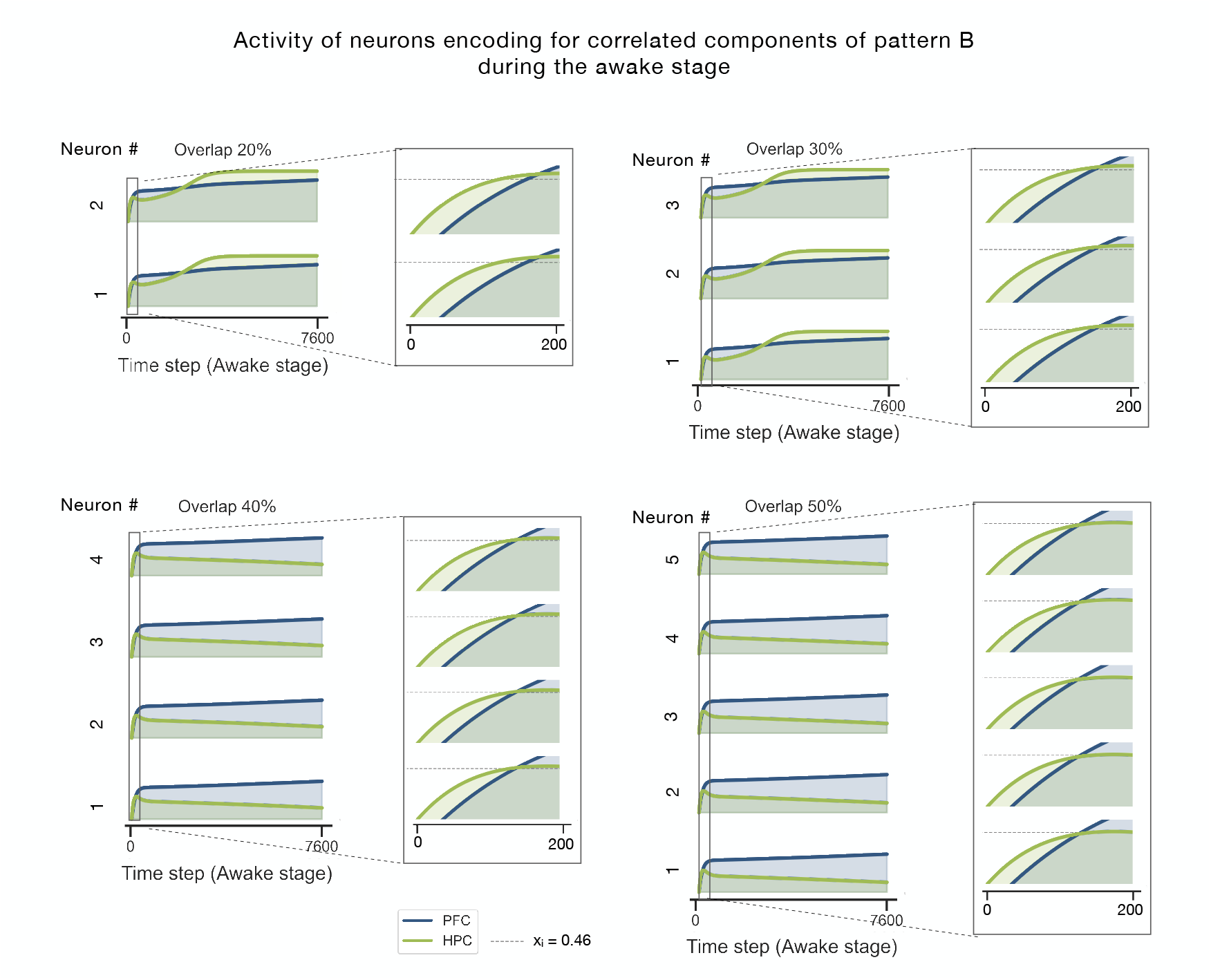
HPC and PFC activity of neurons encoding for correlated components of a pattern B overlapping by 20, 30, 40 and 50% with A. If overlap is 20 or 30%, by the time the HPC neurons encoding for the correlated components of pattern B with A reach an activation level of 0.45, the activity of the corresponding PFC neurons is smaller, and therefore they are not inhibited by PFC. On the other hand, if the overlap is 40 or 50%, the PFC neurons will get to a state where their activity is bigger than the HPC before the later reaching 0.45, and suppress the activity of the HPC neurons encoding for the correlated components. This happens because if the overlap between pattern B and the stored pattern A is big enough, sufficient recurrently connected PFC neurons (i.e. neurons encoding for the correlated components of the patterns) are going to be recruited, resulting in a faster increase in PFC neural activity.

**Figure S5:**
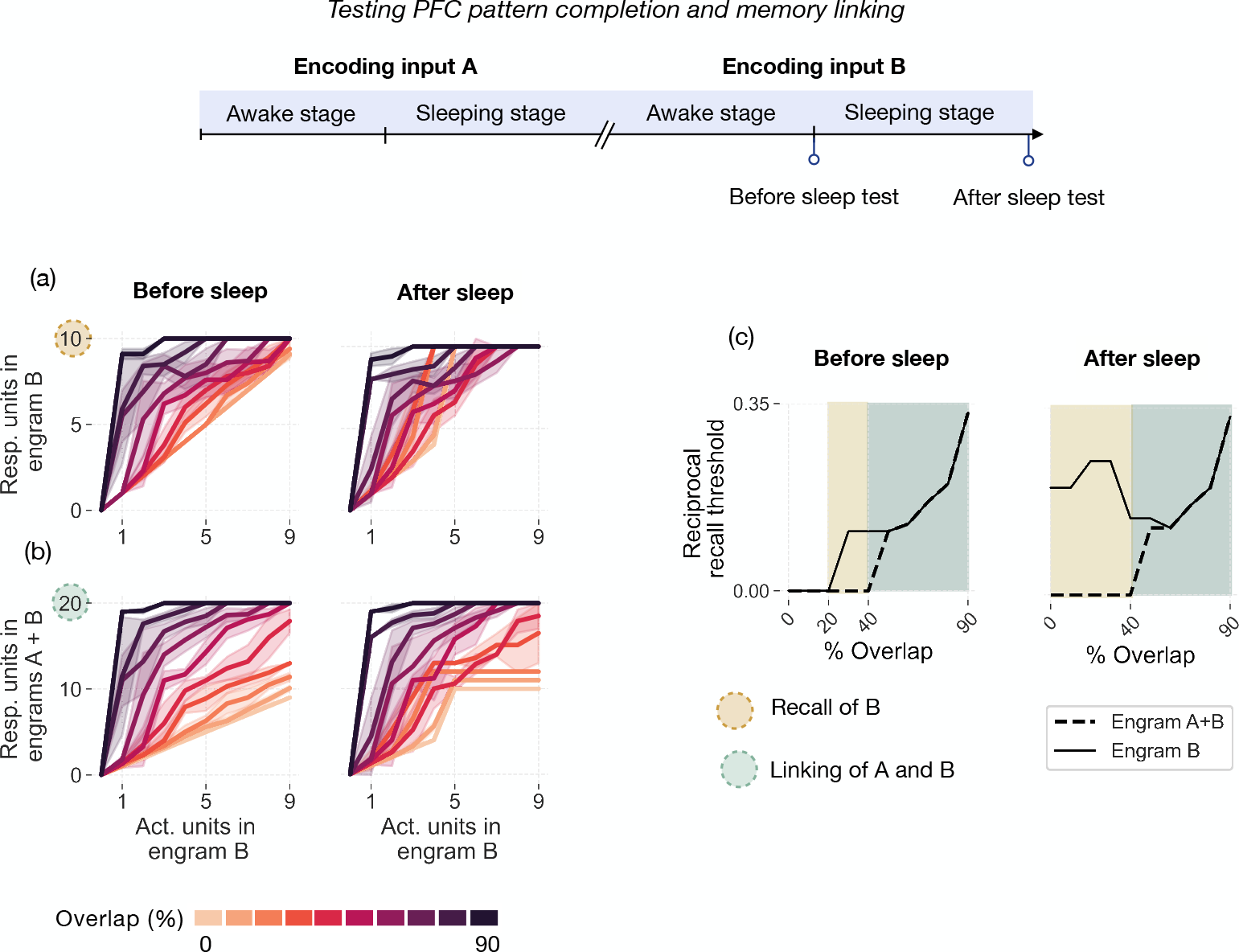
Testing PFC pattern separation and linking of different patterns overlapping from 0-90%. **(a)** Evaluating the PFC ability to recall engram B at the end of the awake stage (before sleep test) and the sleeping stage (after sleep test). At the end of each stage, we activated subsets of engram B neurons of different sizes (ranging from 1 out of 10 to 9 out of 10 engram B neurons) and measured the number of active engram B cells after 7000 time steps. We repeated this process 50 times for each subset size, randomly selecting the engram B cells to form the subset. If 10 out of 10 engram B cells are responsive, the PFC has performed pattern completion of engram At the end of the awake stage, only familiar inputs (overlap *≥* 30%) can be recalled (left panel), whereas at the end of the sleeping stage, pattern B could be recalled independently of its degree of overlap with memory A (right panel). **(b)** Evaluating linking of patterns A and B in PFC before sleep (left panel) and after sleep (right panel). At the end of the awake stage, linking of patterns A and B occurred for familiar inputs (overlap *≥* 40%). In other words, activation of an engram B subset can activate all 20 neurons forming engram A and engram B. At the end of the sleeping stage, inputs with an overlap of *≥* 40% were linked in PFC. **(c)** The minimum number of cells required to successfully perform pattern completion, which we refer to as the “Recall threshold,” was determined based on the minimum number of engram B cells needed to be activated to get all 10 engram B neurons (recall B) or all 20 engram A and B neurons (memory linking) responsive. For example, before sleep, if pattern B overlaps by 40% with memory A, we need to activate 9 out of 10 neurons to get 10 responsive engram B neurons. This means that the recall threshold for input B is 9 (and the reciprocal recall threshold 1/9 *≈* 0.1). A reciprocal recall threshold of zero means that it is not possible to recall the respective engrams. Pattern separation is identified as the cases for which we have recall of B without linking of A+B.

**Figure S6:**
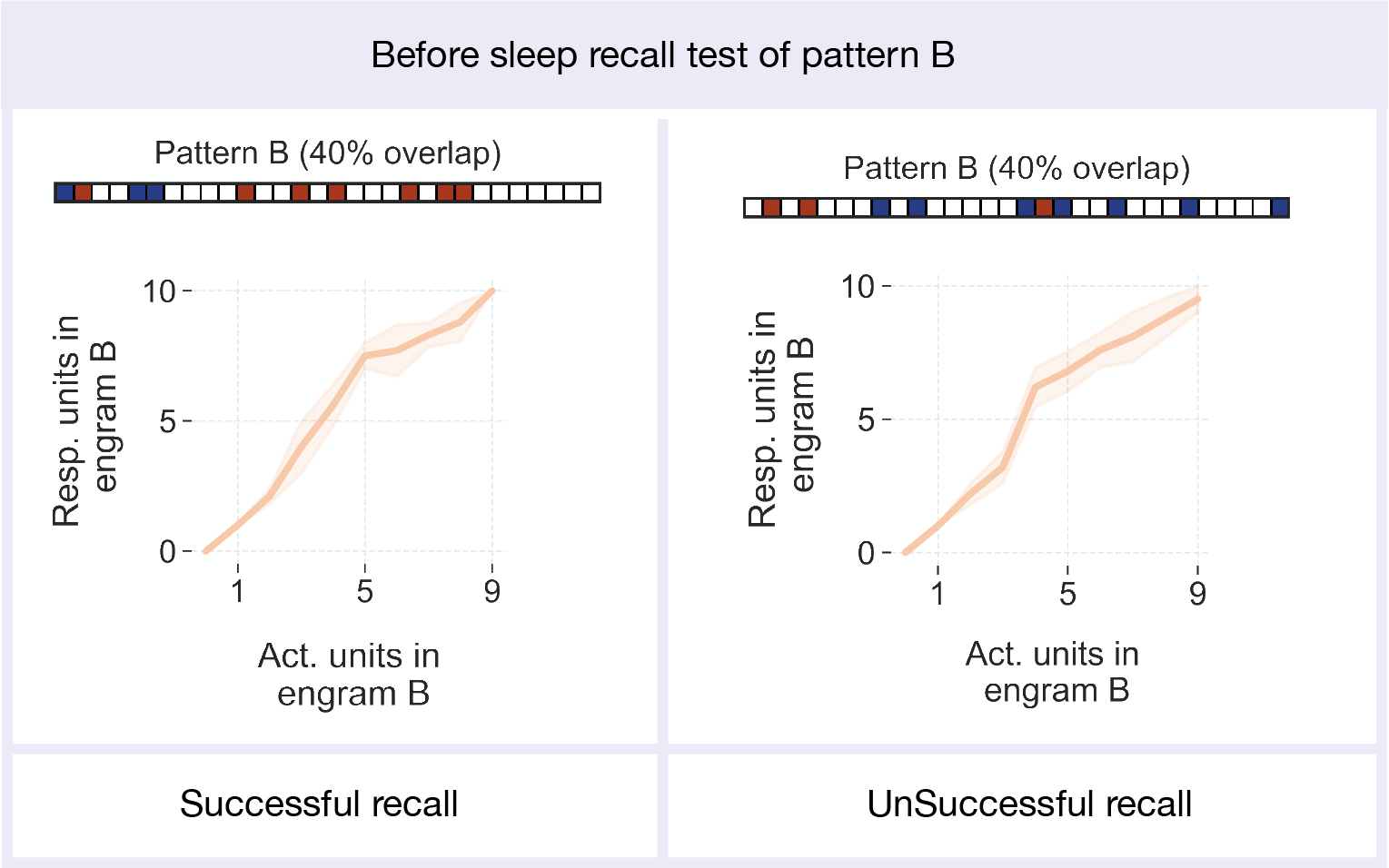
Testing recall of two different patterns B overlapping by 40% with A at the end of the awake stage. Despite having the same degree if similarity, the strength of consolidation in PFC (quantified in the same way as in figure S5) of both patterns differs at the end of the awake stage. Right panel: we can recall pattern B by activating 9 out of 10 PFC neurons encoding it. Left panel: we cannot recall pattern B, regardless of how many PFC neurons encoding memory B we activate.

## Notes

### Competing Interest Statement

The authors have declared no competing interest.

